# Dynamic applicability domain (*d*AD) for compound-target binding affinity prediction task with confidence guarantees

**DOI:** 10.1101/2022.08.22.504786

**Authors:** Davor Oršolić, Tomislav Šmuc

## Abstract

Increasing efforts are being made in the field of machine learning to advance the learning of robust and accurate models from experimentally measured data and enable more efficient drug discovery processes. The prediction of binding affinity is one of the most frequent tasks of compound bioactivity modelling. Learned models for binding affinity prediction are assessed by their average performance on unseen samples, but point predictions are typically not provided with a rigorous confidence assessment. Approaches such as conformal predictor framework equip conventional models with more rigorous assessment of confidence for individual point predictions. In this paper, we extend the inductive conformal prediction (ICP) framework for the dyadic data, such as compound-target binding affinity prediction task. The new framework is based on dynamically defined calibration sets that are specific for each testing interaction pair and provides prediction assessment in the context of calibration pairs from its compound-target neighbourhood, enabling improved guarantees based on local properties of the prediction model. The effectiveness of the approach is benchmarked on several publicly available datasets and through testing in more realistic scenarios with increasing levels of difficulty on a bespoke, complex compound-target binding affinity space. We demonstrate that in such scenarios, novel approach combining applicability domain paradigm with conformal prediction framework, produces superior confidence assessment with informative prediction regions compared to other *state-of-the-art* conformal prediction approaches.

## 1 Introduction

The fast growth of experimental results published in the scientific literature and data available through repositories makes modelling of binding affinity between compounds and protein targets expanding and interesting from both scientific and industrial aspects. Methods for *in silico* modelling and screening large chemical compound spaces are often computational pipelines based on feature generation tools and machine learning algorithms Cichonska et al. [2017] Cichońska et al. [2021] Öztürk et al. [2018] Nguyen et al. [2021]. Compound and target spaces can be described with a multitude of different descriptors, including those that describe their structural or physicochemical properties Lim et al. [2021]. More recent, deep learning algorithms are capable of simultaneously learning specific representations of the input data and the predictive model Öztürk et al. [2018] Nguyen et al. [2021]. For example, convolutional neural networks have found application in both biotechnology and biomedicine Chen et al. [2016] Öztürk et al. [2018] Nguyen et al. [2021]. In Öztürk et al. [2018] they used convolutional neural networks to learn small molecule and protein representations from 1-dimensional sequences. On the other hand, to improve the predictive power of the model with a more realistic representation of molecules, the GraphDTA method was proposed in Nguyen et al. [2021]. This approach is based on a graph convolutional block that learns compound representations from a molecular graph in which atoms are represented as nodes and connections between atoms as graph edges Kipf and Welling [2016]. Each node is characterised by the corresponding atom related features (i.e. the number of adjacent atoms, the number of adjacent hydrogen atoms, etc.) Nguyen et al. [2021] Ramsundar et al. [2019]. Furthermore, protein sequences were represented as finite length integer sequences, with every amino acid denoted as an integer based on an alphabetical symbol Nguyen et al. [2021]. In QSAR modelling predictive models should not only aim to achieve high accuracy on unseen samples, but it is also important that these predictions are accompanied with some confidence guarantees. Conventional QSAR modelling uses applicability domain (AD) to improve prediction credibility Gadaleta et al. [2016]. AD of the QSAR model represents a bounded chemical space within which the model is guaranteed a well-defined and reliable level of average performance Aniceto et al. [2016] Klingspohn et al. [2017] Mathea et al. [2016]. However, the concept of AD does not provide an apparatus that would determine how reliable certain model predictions are Aniceto et al. [2016]. When using AD, the user determines the portion of external data that falls within the established boundaries, without assessing the AD boundary’s ability to differentiate between ‘acceptable’ and ‘unacceptable’ new predictions Aniceto et al. [2016] Mathea et al. [2016]. Intuitively, the applicability domain increases the confidence of the model’s predictions, but this is not directly quantified. According to Aniceto et al. [2016], AD boundaries are set using training sample similarity thresholds or class probability estimates, etc. Klingspohn et al. [2017]. This approach treats AD as a space between the defined limits typically overlooking the possibility of localised holes in the chemical space where the model’s predictions may be unreliable. A well-balanced AD definition would need to include information on distribution of both entities, with separately defined boundaries, when dealing with binding affinity between two different entities (dyadic data).

Conformal prediction (CP) framework was introduced by Gammerman et al. [2013], with the intention of providing confidence guarantees for classification predictions made by the support vector machines. First attempt at conformal prediction was made using transductive conformal predictors (TCP), which required retraining of the model for each individual prediction and were therefore computationally expensive Shafer and Vovk [2008]. As a result, the inductive conformal prediction (ICP) framework was developed Papadopoulos [2008]. The ICP framework uses a calibration set, that typically accounts for a smaller portion of the training samples, to calibrate the trained model and compute confidence levels. The disadvantage of taking such an approach is that the prediction regions computed based on the calibration set for any particular level of confidence are fixed, meaning that they do not alter from test sample to test sample Johansson et al. [2014]. In the conformal prediction framework, AD related measures Klingspohn et al. [2017] can also be used for conformity scoring to locate the applicability domain or the calibration set, for each individual test sample. We call this set a conformity region of sample x. The typical nonconformity function that is utilised for regression tasks is the absolute difference between the true label and the predicted label for a particular sample, |*y*_*i*_ *ŷ*_*i*_| Shafer and Vovk [2008], Johansson et al. [2014], Papadopoulos and Haralambous [2010], Papadopoulos et al. [2011]. This approach focuses solely on the nonconformity in the label space, and assumes that training and calibration sets must be exchangeable Shafer and Vovk [2008]. For a predefined confidence level prediction regions are fixed, meaning they remain the same for any new test sample tested which prevents them from being as informative as we would like them to be. In order to alleviate this, a normalisation step is included in the context of nonconformity function definition Papadopoulos and Haralambous [2010]Papadopoulos et al. [2011], i.e. the standard nonconformity scores are divided by the model accuracy estimated at the location of a given test sample x Papadopoulos and Haralambous [2010]. This method assumes an independent approach for estimating the underlying binding affinity prediction model’s accuracy or error Papadopoulos and Haralambous [2010] Papadopoulos et al. [2011].

In this paper we introduce the conformal prediction framework that is better suited for the dyadic data character of the compound-target binding affinity modelling task. We rationalise that binding affinity modelling task, or for that matter any dataset containing interactions between two entities, requires special treatment when defining nonconformity in the space of compound-target pairs. For binding affinity modelling specifically, binding affinities are conditioned on input data that comes from two distinct distributions: one of compounds and one of targets. Conformal predictors based solely on label data distributions cannot capture the true nonconformity of the input data. We thus aim to adjust the conformal prediction framework for this type of problem by introducing the concept of the dynamic calibration set, the calibration set that is specific for the particular compound-target pair being tested. We test our method and compare it against other state of the art approaches proposed in Shafer and Vovk [2008] Papadopoulos and Haralambous [2010] Papadopoulos et al. [2011] over several standard benchmark compound-target binding affinity datasets, as well as specifically designed small compound protein kinase binding affinity dataset. This dataset combines publicly available databases containing small compound bioactivities of diverse sets of compound scaffolds across the human kinome, and is constructed to allow the testing of the conformal predictor frameworks in settings that are representative of the real use-case scenarios.

## 2 Proposed conformal prediction framework

The initial step in a conventional inductive conformal prediction framework is to divide the training set into a proper training set and a calibration set Shafer and Vovk [2008] Alvarsson et al. [2021]. The calibration set must reflect the distribution of the training samples, satisfying the assumption that the data are independent and identically distributed, or a more relaxed assumption that they are exchangeable Papadopoulos et al. [2011] Shafer and Vovk [2008]. In order to avoid reducing the training space and taking one fixed, works-for-all calibration set, we train a model on all training samples and define a dynamic calibration set such that for each test sample pair x (compound-target pair) *k* most similar compounds and *q* most similar protein targets are selected from the training space, with respect to the tested compound-target pair. *k* and *q* are numbers of nearest neighbours in the compound and target space that determine which training compound-target pairs (existing compound-target pairs and their experimentally measured binding affinities) will be selected for a dynamic calibration set for the particular tested pair. Contrary to the conventional calibration set definition, which is stationary and represents the overall training space, we define a dynamic calibration set for each individual test sample by locating the most similar (conforming) samples of the training landscape to the sample that is being tested. Let us denote the training set of compound-target pairs as:

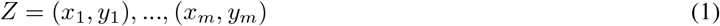

where *x*_*i*_ is a compound-target pair and *y*_*i*_ is a measured binding affinity for that pair. Let the set of compounds and targets be denoted by *C* and *T* respectively, and the corresponding set of training compound-target pairs (tuples) denoted by *X* = (*C, T*). Calibration set (*Z*^*c*^) of a new test sample is (dynamically) constructed from training samples that have maximum Tanimoto similarity coefficients (*s*) towards the tested compound-target pair. Let *x* = (*c, t*) be the new test sample for which we choose a subset of *X* by retrieving *C* ⊂ *C*^*t*^, |*C*| = *k*, such that *s*(*c*^(*i*)^, *c*) ≥ *s*(*c*^(*j*)^, *c*) holds ∀ ∈ *c*^(*i*)^ *C* and ∀*c*^(*j*)^ ∈ *C*^*t*^\*C*. Equivalently, we define *T* ⊂ *T*^*t*^, a subset of targets such that |*T*| = *q. k* and *q* are tunable hyperparameters, determining the neighbourhood of the tested *x* = (*c, t*) pair in the training set X. Dynamic calibration set for the new test instance *x* is then defined as:

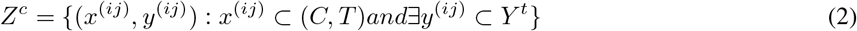

where each *x*^(*ij*)^ is actually a tuple (*c*^(*i*)^, *t*^(*j*)^). The dynamical calibration set defined in this manner represents the most conforming part of the training set bioactivity space with respect to the tested compound-target pair. To allow forming dynamic calibration sets from training samples, we train the model over the entire dataset by applying 10 times repeated 10-fold CV and calculate nonconformity scores for each training sample based on these CV predictions Vovk et al. [2018]. We call the proposed approach of forming dynamic calibration sets - dynamic Applicability Domain (dAD) and define two alternative nonconformity scores which are defined in the following section.

### 2.1 Definition of calibration and test nonconformity scores

In this work we define and test two alternative formulations of nonconformity scores for the compound-target pairs from the dynamic calibration set. The first variant, dAD (NN), nonconformity score 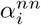 (3) is calculated by taking the difference between the experimental binding affinity of each pair in the *Z*^*c*^ and mean label value for all pairs in *Z*^*c*^. In alternative formulation, dAD (CV), nonconformity score 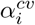 (4) is based on a difference between the experimental binding affinity from the mean of 10×10-fold cross-validation predictions for each pair. For the test sample, the nonconformity scores are defined as the difference of the predicted label (*ŷ*) and each experimental compound-target pair binding affinity in the dynamic calibration set, *α*^*x*^ (5). Thus, for every new test instance *x* we get the corresponding calibration nonconformity vectors, *S*^*cv*^ or *S*^*nn*^ and its own vector of nonconformity scores *S*^*x*^.

Definitions of nonconformity scores associated with the test sample and its dynamic calibration set are given below:

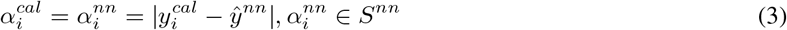

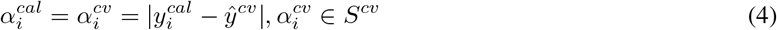

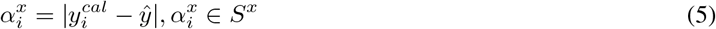

In the next step we have to find the true prediction region for a predefined confidence level for the test sample *x*. For that purpose we have to find minimal value from the calibration nonconformity scores 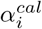, for which the expression below holds Shafer and Vovk [2008]Johansson et al. [2014]:

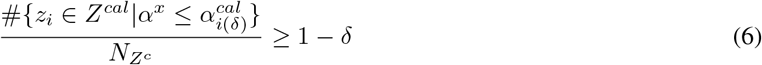

where *z*^(*i*)^ ∈ *Z*^*c*^ are samples from the calibration set. We annotate minimal value of 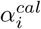 for which above expression holds 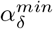. Then the prediction region for any new example is defined as 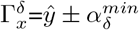. As one can notice, for any predefined level of confidence, the prediction region varies between test samples as nonconformity scores for the test sample are based on the samples in the dynamic calibration set. Chosen 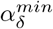 represents the partition of samples in the *S* = *S*^*cv*^ or *S* = *S*^*nn*^ that have higher nonconformity scores than any given 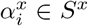. Since tentative labels for each tested interaction pair *x* are based on dynamic calibration set samples, and reflect the local model performance, no normalisation step is performed and individual prediction regions are directly inferred from 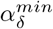.

#### Algorithm 1

dAD (CV/NN)

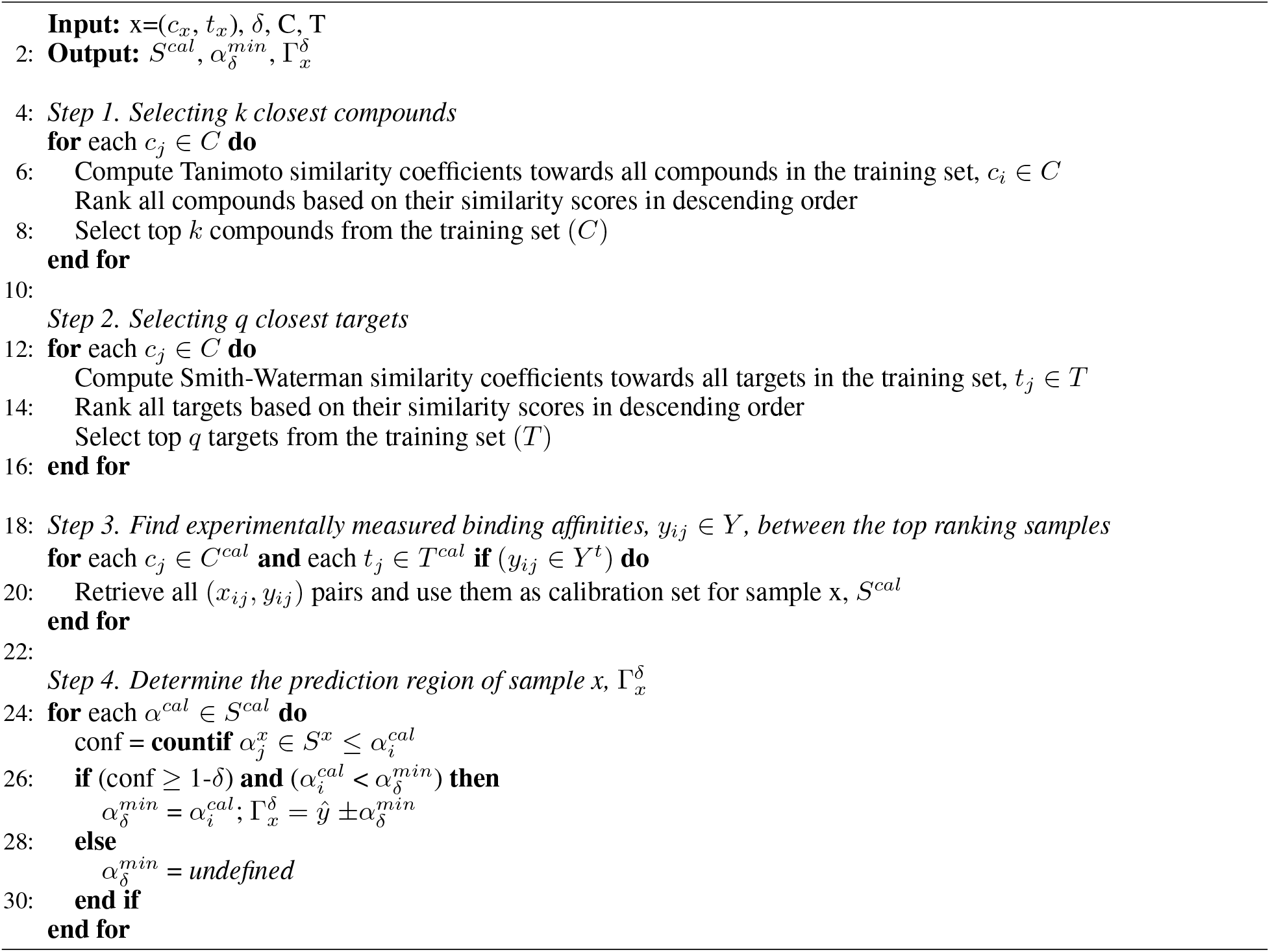

### 2.2 Normalised nonconformity measures

Approaches like Papadopoulos [2008]Johansson et al. [2014]Alvarsson et al. [2021]Papadopoulos et al. [2011], rely on fixed calibration sets and for that reason prediction regions these methods output are fixed and don’t reflect the true nonconformity of test samples where larger prediction regions (or *α*) correspond to non-conforming samples and tighter intervals correspond to well conforming samples. Thus, in order to achieve more informative prediction regions for an individual test sample *x*, studies like Papadopoulos et al. [2011]Papadopoulos and Haralambous [2010] introduced different normalisation measures.

Here we will implement and compare proposed approach with the original paper Shafer and Vovk [2008], and updated versions with normalisation steps introduced in Papadopoulos and Haralambous [2010]Papadopoulos et al. [2011]. In Shafer and Vovk [2008] nonconformity score is calculated simply as the absolute difference of the predicted and the true label:

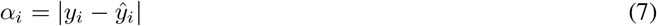

In terms of normalisation, one of the normalisation steps proposed in Papadopoulos and Haralambous [2010] requires building an additional model to estimate the accuracy, or error, for each prediction, *ŷ*_*i*_:

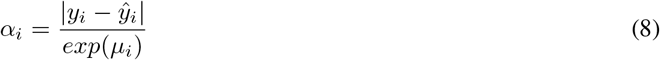

where *μ*_*i*_ represents the natural logarithm of the prediction of absolute error of underlying model. Instead of building a secondary model and predicting errors, normalisation of given *α*_*i*_ can be performed using summation of the distances 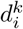 of nearest neighbours of the sample *x*_*i*_ normalised by median of summarised distances of nearest neighbours of each training sample, 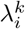, as proposed in Papadopoulos et al. [2011]:

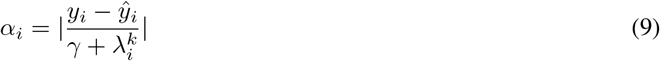

Or, by the standard deviation of true labels for k nearest neighbours of *x*_*i*_ normalised by the median standard deviation of k nearest neighbour labels of every train sample, 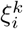, as proposed in Papadopoulos et al. [2011]:

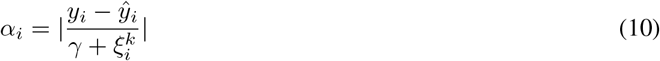

with *γ* being the parameter in control of the sensitivity to changes for both measures 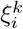 and 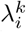 in (9) and (10) Papadopoulos et al. [2011]. Further in this paper we compare the performance of the original study by Shafer and Vovk Shafer and Vovk [2008] and normalised nonconformity scores (8), (10) and (9) with the proposed dAD method, in terms of tightness of outputted prediction regions and their validity.

## 3 Data

For the purpose of determining the validity of the proposed approach, we test it over several publicly available databases as shown in Table 1. Furthermore, we combine all mentioned datasets into a single dataset covering larger space of kinase inhibitors over extensive human kinome space. Aside from the kinase inhibitors, we retrieve the G protein-coupled receptors (GPCR) and selective serotonin re-uptake inhibitor (SSRI) subsets from the DrugTargetCommons Tang et al. [2018] database, in order to test the performance across more diverse bioactivity spaces.

**Table 1:**
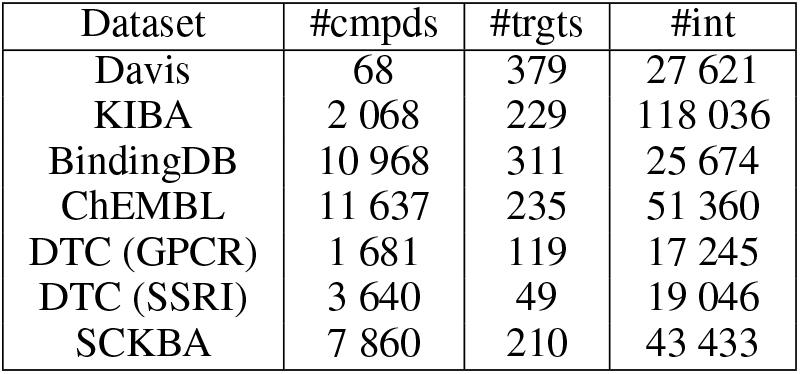
Databases used for compound-target binding affinity prediction, with Davis Davis et al. [2011], KIBA Tang et al. [2014], BindingDB Gilson et al. [2016], ChEMBL Gaulton et al. [2012] and DTC Tang et al. [2018] being publicly available and the small compound-kinase binding affinity (SCKBA) dataset we constructed from the mentioned databases for the purposes of a unified representation of this specific bioactivity space.

Data preprocessing involved several preprocessing filters to ensure that the final datasets contained only those bioactivity profiles that optimally represent the problem. This was accomplished by removing duplicates from benchmarks such as Davis and KIBA datasets, and ensuring that the final version of ChEMBL and BindingDB datasets contained only those bioactivity profiles measured over the human kinome superfamily narrowing the potential protein target space down to 9 distinct kinase groups.

Acquiring larger kinase inhibitor dataset by combining available databases makes it possible to construct several testing scenarios. To construct a more consistent dataset and a well-rounded representation of the bioactivity space we apply several filters to make sure the final dataset contains only protein targets belonging to the human kinome and small compounds with a molecular weight of less or equal to 900 Da. To increase the number of measured bioactivities we use both K_*d*_ and *K*_*i*_ interchangeably, as it was shown in Cichońska et al. [2021] that combination of bioactivity types can increase model’s overall performance. Finally, to reflect the real machine learning use-cases with increasing levels of difficulty, the data was distributed between the training set and four different test sets, as it was introduced in Cichonska et al. [2017] Papadopoulos [2008], Figure 1. For the construction of testing sets in this manner, we used t-SNE and performed chemical space analysis of 7860 compounds (Figure 1, Supplementary). Training compounds with their bioactivity profiles were collected to assure high coverage in versatility of compound scaffolds available in the overall compound set, same as compounds from the densest regions of chemical space. For second test set we retrieve compounds from high density regions, mostly including compounds with fewer bioactivity profiles, but with many similar compounds in the training set - while for third test set, bioactivity profiles were chosen based on the kinome space with few experimentally measured bioactivities in the overall dataset. In the fourth scenario, test set contains both compounds and targets that are not represented in the training set. We used these four distinct test scenarios to test the ability of trained models and different conformal predictor frameworks to generalise over different scenarios representing practical use-cases in the drug-discovery context.

**Figure 1:**
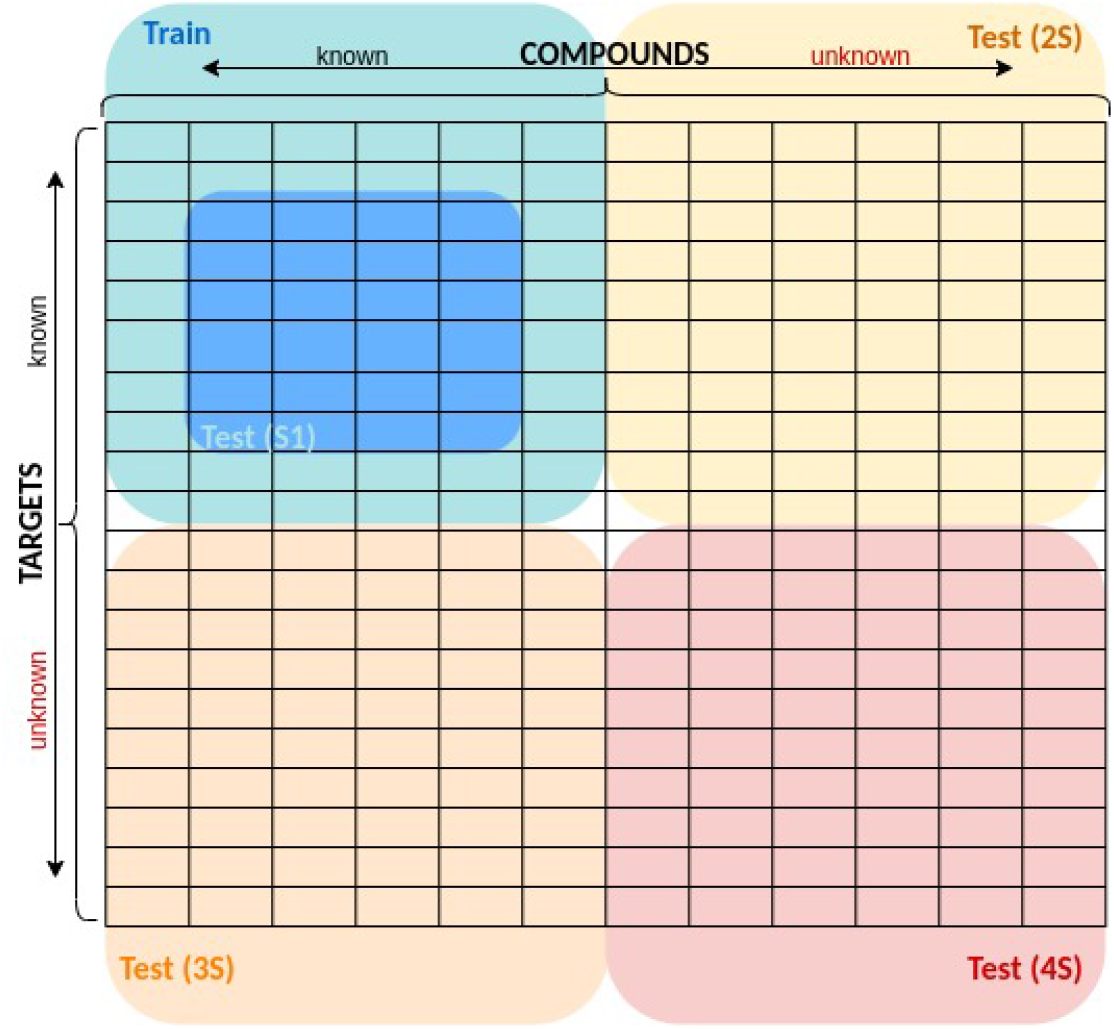
Schematic illustration of test set construction with 4 difficulty levels, where **S1** contains new compound-target pairs and reflects a standard way of testing and evaluating model performance by using stratified sampling and making predictions over known compounds and targets; **S2** contains new compound-target pairs with compounds not available in the training set; **S3** contains new compound-target pairs with targets not available in the training set; and **S4** contains never seen compounds nor targets.

### 3.1 Compound space representation

We build predictive models based on two distinct approaches: first one using XGBoost algorithm and Tanimoto similarity scores calculated using 2048-bit Morgan fingerprints, with radii 2 and applying a pairwise comparison of the whole compound space, and second employing graph convolutional network that learns distributed molecular representation, as a *state-of-the-art* method introduced in Nguyen et al. [2021]. For training GCN on the chemical space, molecules are represented as adjacency matrices between atoms, and each atom is represented as a vector of properties. Instead of using one-hot-encoded vector representation Nguyen et al. [2021], we use *rdkit* library Landrum et al. [2020] in Python to compute atomic attributes that include the atomic number, charge, hybridisation state, number of radical electrons, number of hydrogen atoms bound, chirality, and ring membership. Deep learning approach is implemented using PyTorch Paszke et al. [2019] and PyTorch Geometric He et al. [2022].

### 3.2 Target space representation

To test the new framework in realistic scenarios we use as the chemical space protein kinases, as it is one of the largest protein groups of pharmacological interest Liao [2007]. The human kinome consists of approximately 530 enzymes clustered into 10 smaller groups or super-families that share a common evolutionary origin Roskoski Jr [2015]. Most importantly, they catalyze phosphorylation and are included in most important regulatory mechanisms in all living organisms Roskoski Jr [2015] Roskoski Jr [2014]. Local similarities between these protein targets are computed by applying the Smith-Waterman algorithm to the protein kinase sequences with default parameters of the *protr* library in *R* Xiao et al. [2015] (gap.opening=10, gap.extension=4) and *BLOSUM* 62 substitution matrix. Same approach was performed for computation of sequence similarities of the GPCR and SSRI datasets, retrieved from the DrugTargetCommons database Tang et al. [2018]. Specifically for the SCKBA dataset, we compute local similarities only for the protein kinase domain (PK), because it is highly conserved and is a major focus for the small molecule inhibitor design, with most of the approved drugs targeting exactly ATP-binding cleft or the surrounding regions Liao [2007]. For the training of convolutional neural network architecture (GCN-CCN) on SCKBA dataset, the same strategy is adopted as in Nguyen et al. [2021], treating protein targets as a sequences of characters within a CNN block. The only difference is that instead of using the entire protein sequences, we learn sequence representations by feeding PK domain sequences into the network, for reasons explained above. We identify the longest PK domain sequence in our dataset and pad other targets to match the length of the identified sequence.

## 4 Methods

XGBoost is trained on individual compound-target binding affinity datasets and on a SCKBA dataset, with binding affinities expressed as the negative logarithm of equilibrium of dissociation (*K*_*d*_) or inhibition (*K*_*i*_) constants, see Table 1, while GCN-CNN architecture is applied only for the SCKBA dataset.

### XGBoost

It is a decision tree based ensemble method that has proven to be fast and highly effective in achieving *state-of-the-art* results on many problems Chen and Guestrin [2016]. There are several hyperparameters determining the quality and performance of the XGBoost model for a given task; we utilised grid-search approach to find the set of hyperparameters providing best performing model. Since hyperparameter tuning in this manner can become expensive, smaller subset of the SCKBA dataset was used for this purpose.

### GCN-CNN

For the GCN-CNN approach, we started from GraphDTA architecture proposed in Nguyen et al. [2021]. We updated it by changing node representations of compound graphs from one-hot-encoded vectors to the physicochemical atomic feature representations. Furthermore, NN architecture is customised by implementing early stopping for no significant improvement over 20 consecutive epochs, with the decrease in learning rate for every 10 epochs, in order to avoid over-fitting the model on the training data (Figure 4, Supplementary). GCN-CNN approach is trained for the total of 260 epoch with starting learning rate of 0.005.

### CP

For computing calibration scores for any of the baseline approaches, we use Python library **nonconformist**. As it is shown in Figure 2, in order to asses the validity of the confidence levels (75%, 80%, 85%, 90%, 95%, 99%), the trained model is subjected to four different levels of testing difficulty. Additionally, to prove reliability of this approach we test it on additional datasets from the Table 1. Validity of predictions over realistic use-case scenarios is tested only on the SCKBA dataset, due to the size and complexity of the compound-target space it represents. In order to compare proposed approach with baseline studies, we implement normalised nonconformity scoring as proposed in Papadopoulos and Haralambous [2010] Papadopoulos et al. [2011]. For normalising the nonconformity scores by additional error model, instead of training NN Papadopoulos and Haralambous [2010] or random forest Johansson et al. [2014], in this work we train an XGBoost model with the same hyperparameters as the model trained to predict binding affinities.

**Figure 2:**
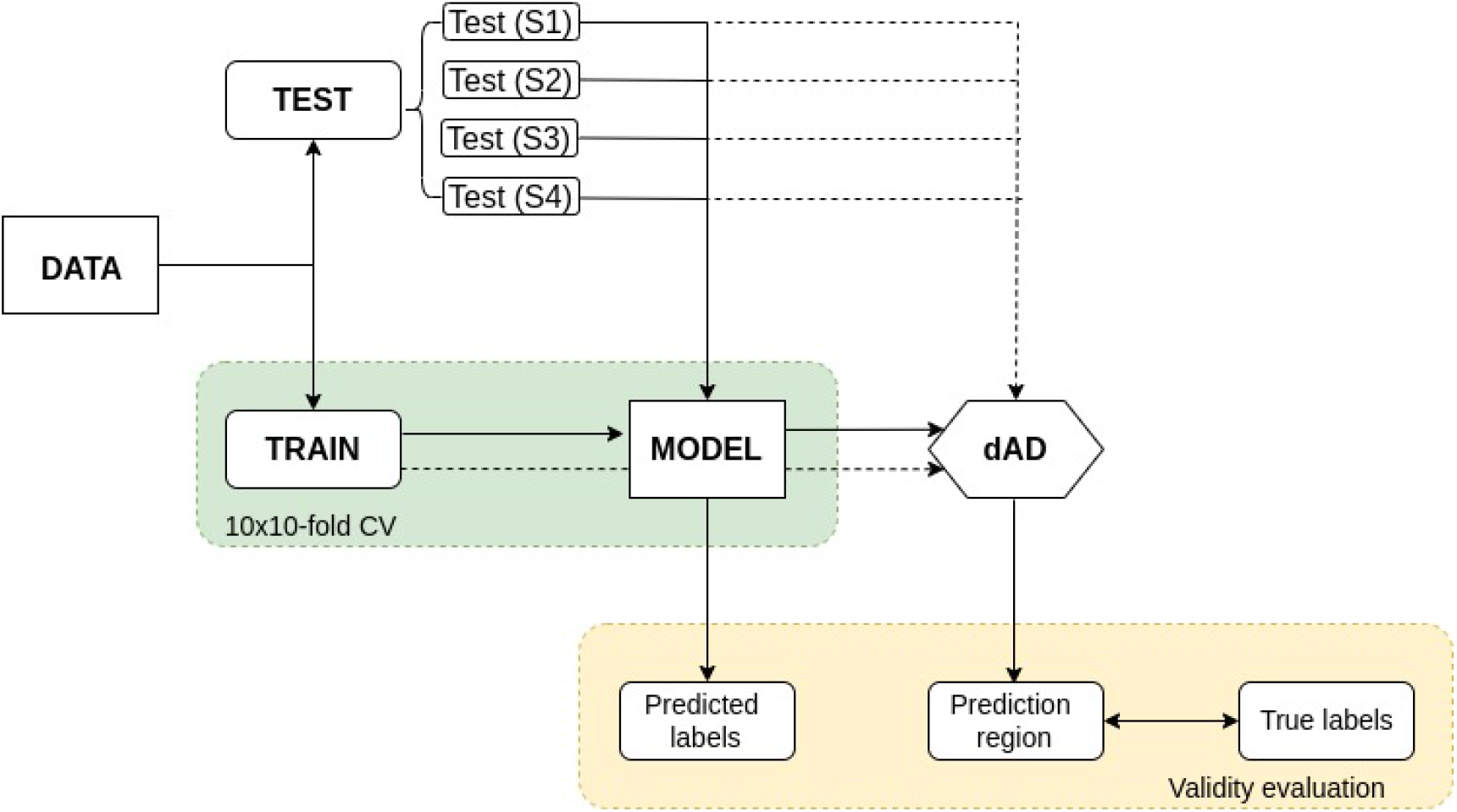
Schematic illustration of the proposed dAD workflow from training of the underlying model to conformal prediction and validity evaluation. Data is split on train and test sets, with all training samples being used for the modelling and allocation of the conformity region for each individual sample in the test set resulting in varying prediction regions. Prediction regions are valid if the true label can be found between its lower and upper bound.

As explained in section 2, the calibration set in our approach is dynamically selected based on an individual compound-target pair tested. We test two variants of nonconformity scores in context of the proposed dAD approach: (i) those based on cross-validation predictions, (dAD-CV) and (ii) using the mean of labels of all nearest neighbours of the tested compound-target pair, (dAD-NN). The dynamic calibration set for any tested sample is based on the existing binding affinities of *k* compounds and *q* targets that are closest in proximity to the compound and protein target from the tested pair *x*. Both hyperparameters, *k* and *q*, were tuned manually for the SCKBA test (S1) dataset, with the aim of inspecting the impact on the validity and size of the prediction regions. However, as it is shown on Figure 5 (Supplementary) for confidence levels of 80%, 90% and 99%, change in the any of the two values did not significantly affect the validity of prediction regions, but it did impact the size of the calibration sets, consequently impacting how informative the output prediction regions are. For the majority of cases with larger number of different compounds and targets we used *k*=250 for nearest neighbours in the compound space and *q*=25 for nearest neighbours in the target space, with the exception when it comes to the Davis Davis et al. [2011] and SSRI Tang et al. [2018] datasets. Davis consists of 68 compounds in total and SSRI consists of only 49 protein targets, so smaller values for *k* and *q* were used, *k*=25 and *q*=10, respectively. Notably, we also tune a *gamma* sensitivity parameter for every dataset individually. Using the narrowest median prediction region (*α*_*δ*_) as a reference, we picked an appropriate value for *γ* under the restriction that the mean error rate does not exceed the mean of the maximum error rates while still retaining the validity of the prediction regions for each confidence level (Figure 6-7, Supplementary). However, the proposed dAD approach due to the dynamic nature of the calibration set construction does not allow the extraction of 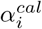 scores for every sample at any confidence level. For that reason, we denote the level of coverage as a value in addition to the reported error rates. Also, for a more direct comparison between the methods, we report the results between them only for those compound-target pairs and confidence levels that our dAD (NN) and dAD (CV) methods were able to produce (Table 1-3, Supplementary).

**Table 2:** XGBoost and GCN-CNN trained on a SCKBA dataset of kinase inhibitors and tested over four distinct difficulty scenarios (S1-S4).

| Train |  |  | Test |  |  |  | XGBoost |  | GCN-CNN |  |
| --- | --- | --- | --- | --- | --- | --- | --- | --- | --- | --- |
| #int | #cmpds | #trgts | SX | #int | #cmpds | #trgts | MSE | CI | MSE | CI |
| 40 578 | 6 325 | 206 | S1 | 655 | 411 | 169 | 0.28 | 0.83 | 0.43 | 0.82 |
|  |  |  | S2 | 935 | 935 | 59 | 0.61 | 0.79 | 0.95 | 0.75 |
|  |  |  | S3 | 655 | 439 | 4 | 0.60 | 0.72 | 0.61 | 0.74 |
|  |  |  | S4 | 600 | 600 | 1 | 3.97 | 0.53 | 3.74 | 0.53 |

**Table 3:** Comparison of baseline methods with proposed dynamic applicability domain (dAD) approach on a combined small compound-kinase binding affinity dataset over four difficulty scenarios, with sensitivity parameters for (9) and (10) in S1 scenario being *λ*=0.7 and *ξ*=0, respectively, and 0 for the rest. The *SX* denotes the testing scenario; *α*_(*δ*)_ is the median prediction region of the test set; *#calib* is number of samples in the calibration set or median number of samples for the dAD method with varying calibration sizes. Error rates represent the percent of samples with labels outside of prediction regions. Values next to the dAD (CV) and dAD (NN) error rates denote the coverage of the test set, with 0 meaning that the proposed approach was not able to produce prediction regions for a given confidence and 1 meaning that it produced a prediction region for every test sample.

| SCKBA |  |  |  |  |  |  |  |  |  |
| --- | --- | --- | --- | --- | --- | --- | --- | --- | --- |
| Approach | SX | Median |  | Error rates per confidence level (%) |  |  |  |  |  |
| | | $\alpha_{\delta}$ | #calib | 75%CI | 80% | 85% | 90% | 95% | 99% |
| Shafer & Vovk (7) | S1 | 0.86 | 4000 | 21.83 | 16.34 | 11.91 | 6.56 | 3.66 | 0.76 |
|  | S2 | 0.86 | 4000 | 40.64 | 35.08 | 30.27 | 21.60 | 12.41 | 2.57 |
|  | S3 | 0.86 | 4000 | 35.49 | 31.73 | 26.02 | 18.95 | 11.43 | 3.91 |
|  | S4 | 0.86 | 4000 | 86.17 | 84.50 | 81.83 | 80.00 | 72.83 | 49.17 |
| Papadopoulos (8) | S1 | 2.41 | 4000 | 7.02 | 4.43 | 3.21 | 2.14 | 1.37 | 0.31 |
|  | S2 | 1.66 | 4000 | 21.93 | 17.86 | 14.65 | 10.80 | 7.17 | 1.07 |
|  | S3 | 2.33 | 4000 | 10.83 | 7.67 | 6.32 | 4.21 | 2.41 | 0.15 |
|  | S4 | 1.47 | 4000 | 78.5 | 73.83 | 68.33 | 59.67 | 41.00 | 7.33 |
| Papadopoulos (9) | S1 | 1.02 | 4000 | 19.69 | 15.73 | 10.84 | 7.33 | 5.50 | 1.68 |
|  | S2 | 1.13 | 4000 | 30.16 | 24.39 | 19.68 | 13.69 | 7.49 | 1.39 |
|  | S3 | 1.55 | 4000 | 22.71 | 18.35 | 14.74 | 10.68 | 5.86 | 1.20 |
|  | S4 | 3.32 | 4000 | 33.83 | 25.17 | 17.17 | 6.33 | 2.67 | 0.00 |
| Papadopoulos (10) | S1 | 0.85 | 4000 | 22.60 | 16.49 | 12.98 | 7.79 | 4.12 | 1.22 |
|  | S2 | 0.55 | 4000 | 43.10 | 37.75 | 32.09 | 25.67 | 17.11 | 5.67 |
|  | S3 | 0.95 | 4000 | 32.18 | 27.37 | 22.71 | 16.54 | 10.53 | 3.01 |
|  | S4 | 0.78 | 4000 | 87.83 | 85.83 | 84.33 | 81.50 | 76.5 | 55.17 |
| dAD (NN) | S1 | 1.85 | 315 | 1.87 (.73) | 1.65 (.74) | 0.79 (.77) | 1.00 (.76) | 0.62 (.74) | 0.00 (.63) |
|  | S2 | 1.78 | 261 | 8.76 (.74) | 6.72 (.70) | 3.54 (.63) | 2.94 (.58) | 1.28 (.59) | 0.36 (.60) |
|  | S3 | 1.65 | 268 | 11.90 (.69) | 10.11 (.71) | 8.07 (.73) | 6.03 (.77) | 4.76 (.79) | 1.03 (.73) |
|  | S4 | 1.79 | 259 | 70.70 (.26) | 68.40 (.38) | 60.47 (.56) | 52.27 (.84) | 39.22 (.98) | 12.69 (.87) |
| dAD (CV) | S1 | 1.61 | 315 | 1.73 (.62) | 1.79 (.60) | 1.08 (.56) | 1.23 (.50) | 0.42 (.36) | 0.00 (.20) |
|  | S2 | 1.60 | 266 | 9.31 (.61) | 8.01 (.57) | 4.75 (.50) | 3.23 (.46) | 1.34 (.40) | 0.34 (.31) |
|  | S3 | 1.43 | 268 | 12.71 (.64) | 12.02 (.63) | 9.74 (.59) | 7.65 (.55) | 3.81 (.43) | 0.00 (.21) |
|  | S4 | 1.69 | 259 | 70.67 (.25) | 69.20 (.37) | 60.55 (.55) | 52.3 (.76) | 45.41 (.69) | 10.42 (.40) |

## 5 Results and Discussion

We compare the proposed approach with the original CP method Shafer and Vovk [2008] and three normalisation based approaches proposed in Papadopoulos and Haralambous [2010]Papadopoulos et al. [2011], all showing good results in previous studies. To provide a more thorough comparison, each method is tested over a few benchmark datasets containing bioactivity data, and a combined dataset specifically representing small compound-kinase binding affinities.

### 5.1 XGBoost vs. GCN-CNN

We tested two different algorithms and problem representations to produce prediction models for testing and comparison of CP approaches over all datasets. As a conventional and faster approach, XGBoost was used as a baseline method - additionally, we adapt and apply GCN-CNN architecture as a *state-of-the-art* deep representation learning approach. We start with the architecture from Nguyen et al. [2021], with the main difference of choosing more meaningful node representations for the compounds, while instead of taking whole protein sequences, we focused specifically on protein kinase domains comprising of catalytic and regulatory subunits of kinase machinery Roskoski Jr [2015]. If we check the Table 3, we can see that XGBoost approach seems to be comparable, if not better in performance than the more complex convolutional network. This outcome may be unique to this dataset, where extended circular fingerprints capture chemical structure variation in the compound space sufficiently well for the model to generalise to real-world scenarios (S2, S3).

### 5.2 Comparison over different difficulty scenarios

To test how well the conformal regressor can recognise true binding affinities and produce valid prediction regions, the share of falsely classified compound-target pairs is determined for each confidence level of the four test scenarios in SCKBA dataset. As can be seen in Table 3, the dAD approach exhibits lower error rates per confidence level, in comparison to the standard approaches. The performance gap is most pronounced in the testing scenarios S2 and S3, which, respectively, include compounds and targets that were not seen during the training phase. In the testing scenario S1, all approaches seem to be equally effective in keeping the number of incorrectly classified samples within proper limits. In contrast, in the testing scenario S4, neither approach is effective, and all of them show a high number of incorrectly classified samples.

In Figure 4 depicts how median of the prediction region (*α*) varies for different CP approaches; Shafer and Vovk (7) method is represented simply as a point, due to the fact that prediction region is constant for all test samples at given confidence level. Median prediction regions for dAD (NN) and dAD (CV) are comparable, and range from one to three units depending on the confidence level. Variation across methods, in terms of prediction regions remains similar even when only dAD covered test samples are taken into account (Figure 2, Supplementary). However, we cannot make the assumption that one method is superior to another simply by looking at the error rates or the validity of a conformal predictor; but rather also consider the width of the prediction regions for each of these test cases. A good conformal predictor should strike a balance between both of these measures, staying valid while ensuring that prediction regions are as narrow as possible. Large prediction regions, while ensuring validity for any test scenario, are not useful. It is well illustrated in Figure 3A (and Figure 8A, Supplementary), where all six CP approaches are compared by examining the relation of the mean error rates with the median *α*_*δ*_ scores paired and unpaired datasets. Results of both dAD (NN) and dAD (CV) seem to be comparable. Similar case is with the Shafer and Vovk approach and Papadopoulos (10), showing somewhat tighter prediction regions but higher error rates on average than the proposed dAD approach. In Figure 3A we visualise performance of all six methods over four different testing scenarios (S1-S4) by plotting the relationship between the mean error rates and median prediction region width (*α*). For S1 test setting, denoted as dots in Figure 3A, all methods besides Papadopoulos (8) in red, give reasonably tight prediction regions with maximum median prediction region lower than 2 units and with low error rates, which is to be expected due to the nature of that testing scenario comprising of compound and protein kinase targets already seen during the training phase. At the other end of the spectrum, there is an S4 test setting where all trained models show almost random performance, as we see in Figure 3A where the performance over the S4 test set marked as pluses occupies the right end of the scatter plot. Papadopoulos (9) is an exception, achieving very low mean error rate on the forth scenario due to the very wide median prediction region. Both proposed dAD approaches show low mean error rates with relatively tight prediction regions, which is especially important for more realistic test scenarios, S2 and S3.

**Figure 3:**
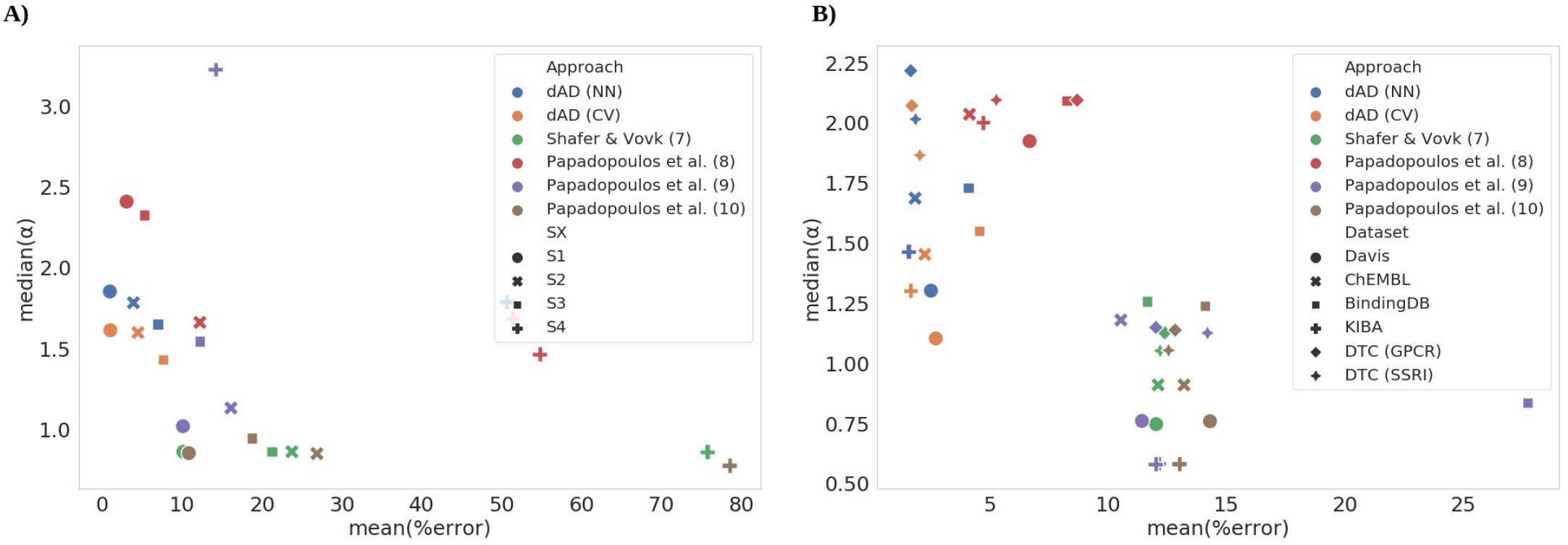
Comparison of the proposed dAD approach with the baseline studies by showing the relationship of mean (%) error and median nonconformity scores for A) four different testing scenarios (S1-S4) of SCKBA dataset and B) over six compound target datasets, Davis Davis et al. [2011], KIBA Tang et al. [2018], ChEMBL database Gaulton et al. [2012], BindingDB Gilson et al. [2016], DTC (GPCR) Tang et al. [2018] and DTC (SSRI) Tang et al. [2018] represented with different shapes.

### 5.3 Comparison over benchmark datasets

To further test the effectiveness of the proposed method, both dAD approaches are evaluated and compared using models trained over several publicly available and frequently used compound-target binding affinity datasets. Furthermore, two sets that have been constructed from the DrugTargetCommons database Tang et al. [2018], GPCR and SSRI inhibitor datasets, Table 1. The Tables 4-5 compare the four conformal predictor approaches with two proposed dynamic approaches on these datasets. Shafer and Vovk approach Shafer and Vovk [2008] shows lower validity of prediction regions when compared to the normalisation based CPs or the proposed approaches. This behaviour is expected, especially since it produces fixed prediction regions (*α* scores) given the confidence level and thus cannot capture individual nonconformity of the test samples. Having this in mind, even if the mean of the prediction region is lower than in any other approach, true difference is shown in the validity of their predictions.

**Table 4:** Comparison of baseline methods with proposed dynamic applicability domain (dAD) approach on compound-kinase binding affinity datasets from Table 1, with sensitivity parameters of equations (9) and (10) for the Davis and KIBA datasets being *λ*=0.7 and *ξ*=0; for BindingDB dataset *λ*=0 and *ξ*=0; and for the ChEMBL dataset *λ*=0.3 and *ξ*=0. Column definitions are the same as in Table 3.

| Benchmark datasets (KI) |  |  |  |  |  |  |  |  |  |
| --- | --- | --- | --- | --- | --- | --- | --- | --- | --- |
| Dataset | Approach | Median |  | Error rates per confidence level (%) |  |  |  |  |  |
| | | $\alpha_\delta$ | #calib | 75% | 80% | 85% | 90% | 95% | 99% |
| Davis | Shafer & Vovk (7) | 0.75 | 1500 | 23.86 | 19.28 | 14.43 | 9.77 | 4.13 | 0.76 |
|  | Papadopoulos (8) | 1.92 | 1500 | 13.12 | 10.36 | 7.63 | 5.32 | 2.76 | 0.91 |
|  | Papadopoulos (9) | 0.76 | 1500 | 23.13 | 18.46 | 13.61 | 9.03 | 3.73 | 0.65 |
|  | Papadopoulos (10) | 0.76 | 1500 | 26.08 | 21.77 | 17.61 | 12.78 | 6.28 | 1.36 |
|  | dAD (CV) | 1.10 | 502 | 3.56 (.34) | 3.33 (.52) | 3.30 (.71) | 3.04 (0.80) | 2.21 (.77) | 0.87 (.46) |
|  | dAD (NN) | 1.30 | 502 | 3.43 (.33) | 3.25 (.52) | 3.24 (.71) | 2.81 (.83) | 1.87 (.91) | 0.48 (.91) |
| KIBA | Shafer & Vovk (7) | 0.58 | 3000 | 23.96 | 19.08 | 14.73 | 9.41 | 4.79 | 0.94 |
|  | Papadopoulos (8) | 2.00 | 3000 | 8.65 | 6.92 | 5.55 | 3.82 | 2.34 | 1 |
|  | Papadopoulos (9) | 0.58 | 3000 | 23.62 | 18.96 | 14.51 | 9.42 | 4.66 | 0.91 |
|  | Papadopoulos (10) | 0.58 | 3000 | 24.85 | 20.01 | 15.73 | 10.23 | 5.77 | 1.46 |
|  | dAD (CV) | 1.30 | 1661 | 3.77 (.73) | 2.62 (0.80) | 1.99 (.87) | 1.04 (.91) | 0.39 (.84) | 0.09 (46) |
|  | dAD (NN) | 1.47 | 1661 | 3.60 (.72) | 2.46 (.81) | 1.85 (.90) | 0.96 (.96) | 0.39 (.98) | 0.1 (.93) |
| BindingDB | Shafer & Vovk (7) | 1.26 | 3000 | 23.36 | 28.85 | 13.08 | 8.84 | 5.00 | 0.84 |
|  | Papadopoulos (8) | 2.09 | 3000 | 13.48 | 11.37 | 8.84 | 6.9 | 5.13 | 3.85 |
|  | Papadopoulos (9) | 0.84 | 3000 | 37.41 | 34.79 | 31.12 | 27.28 | 23.01 | 12.88 |
|  | Papadopoulos (10) | 1.24 | 3000 | 25.52 | 21.6 | 16.42 | 11.78 | 7.45 | 1.83 |
|  | dAD (CV) | 1.55 | 133 | 9.06 (.58) | 6.72 (.55) | 5.45 (.49) | 3.54 (.44) | 1.86 (.39) | 0.80 (.27) |
|  | dAD (NN) | 1.33 | 133 | 8.64 (.73) | 6.08 (.71) | 4.58 (.68) | 3.09 (.67) | 1.54 (.65) | 0.61 (.48) |
| ChEMBL | Shafer & Vovk (7) | 0.91 | 3000 | 24.62 | 19.27 | 13.92 | 9.40 | 4.51 | 0.99 |
|  | Papadopoulos (8) | 2.04 | 3000 | 8.01 | 6.43 | 4.62 | 3.28 | 1.83 | 0.69 |
|  | Papadopoulos (9) | 1.18 | 3000 | 19.79 | 16.01 | 12.13 | 8.77 | 4.91 | 1.66 |
|  | Papadopoulos (10) | 0.91 | 3000 | 25.41 | 20.49 | 15.13 | 10.63 | 5.86 | 1.77 |
|  | dAD (CV) | 1.45 | 253 | 5.18 (.74) | 3.70 (.68) | 2.35 (.62) | 1.47 (.53) | 0.62 (.40) | 0.00 (.15) |
|  | dAD (NN) | 1.69 | 253 | 4.38 (.86) | 3.11 (.86) | 1.85 (.86) | 1.13 (.86) | 0.43 (.83) | 0.15 (.57) |

**Table 5:** Comparison of baseline methods with proposed dynamic applicability domain (dAD) approach on DTC (GPCR) and DTC (SSRI) datasets from Table 1, with sensitivity parameters for (9) and (10) for GPCR dataset being *λ*=0.7 and *ξ*=0; and for SSRI dataset *λ*=0 and *ξ*=0. Column definitions are the same as in Table 3.

| DTC (GPCR; SSRI) |  |  |  |  |  |  |  |  |  |
| --- | --- | --- | --- | --- | --- | --- | --- | --- | --- |
| Dataset | Approach | Median |  | Error rates per confidence level (%) |  |  |  |  |  |
| | | $\alpha_\delta$ | #calib | 75% | 80% | 85% | 90% | 95% | 99% |
| GPCR | Shafer & Vovk (7) | 1.13 | 1500 | 25.02 | 18.85 | 14.29 | 10.00 | 5.16 | 1.07 |
|  | Papadopoulos (8) | 2.09 | 1500 | 13.62 | 11.24 | 9.02 | 7.46 | 6.02 | 4.82 |
|  | Papadopoulos (9) | 1.15 | 1500 | 24.15 | 18.53 | 14.29 | 9.19 | 5.13 | 0.87 |
|  | Papadopoulos (10) | 1.14 | 1500 | 25.28 | 18.93 | 14.93 | 10.44 | 6.18 | 1.28 |
|  | dAD (CV) | 2.14 | 874 | 3.69 (.84) | 2.98 (.81) | 1.84 (.77) | 1.12 (.70) | 0.54 (.59) | 0.1 (.58) |
|  | dAD (NN) | 2.25 | 874 | 3.72 (.94) | 2.82 (.93) | 1.77 (.90) | 1.09 (.85) | 0.45 (.78) | 0.08 (.75) |
| SSRI | Shafer & Vovk (7) | 1.05 | 1500 | 24.7 | 19.13 | 14.59 | 9.53 | 4.25 | 1.05 |
|  | Papadopoulos (8) | 2.09 | 1500 | 10.21 | 8.17 | 6.15 | 3.83 | 2.06 | 1.24 |
|  | Papadopoulos (9) | 1.13 | 1500 | 25.83 | 21.09 | 17.26 | 12.06 | 6.6 | 2.41 |
|  | Papadopoulos (10) | 1.05 | 1500 | 24.57 | 19.69 | 15.33 | 9.87 | 4.7 | 1.21 |
|  | dAD (CV) | 1.86 | 234 | 4.17 (.68) | 3.15 (.63) | 4.47 (.53) | 1.56 (.47) | 0.66 (.40) | 0.23 (.21) |
|  | dAD (NN) | 2.02 | 234 | 3.98 (.89) | 2.99 (.85) | 2.09 (.81) | 1.38 (.74) | 0.63 (.67) | 0.11 (.49) |

On the other hand, normalised approach using the underlying error model and 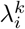 coefficient gives comparable results to the proposed dAD method for 95% and 99% confidence levels on the Davis dataset. The difference is better depicted in boxplots Figure 5 showing the distribution of prediction regions for every *x* in the test set. Results for the same test sets, but averaged only over samples for which dAD methods were capable to define prediction regions are given in Figure 3 in Supplementary. True difference in the performance of any of mentioned methods is shown by scatter plot giving relation of the mean percentage of wrongly classified samples to the median *α* score, Figure 3B (and Figure 8, Supplementary). The ideal conformal predictor would have low error rate (be valid) and have prediction regions as tight as possible, i.e. in this figure the better performing CPs should be closer to zero on both axes. When compared in this sense, approaches (7), (9) and (10) have the narrowest prediction regions when tested on all six datasets, consequently with higher mean error rates ranging above 10%. When taking into consideration both the prediction regions and mean error rates, dAD (NN) and dAD (CV) produce more optimal prediction regions to the error rate relation, Figure 3B, with median prediction regions exceeding the two units only for the DTC (GPCR) and DTC (SSRI) datasets.

**Figure 4:**
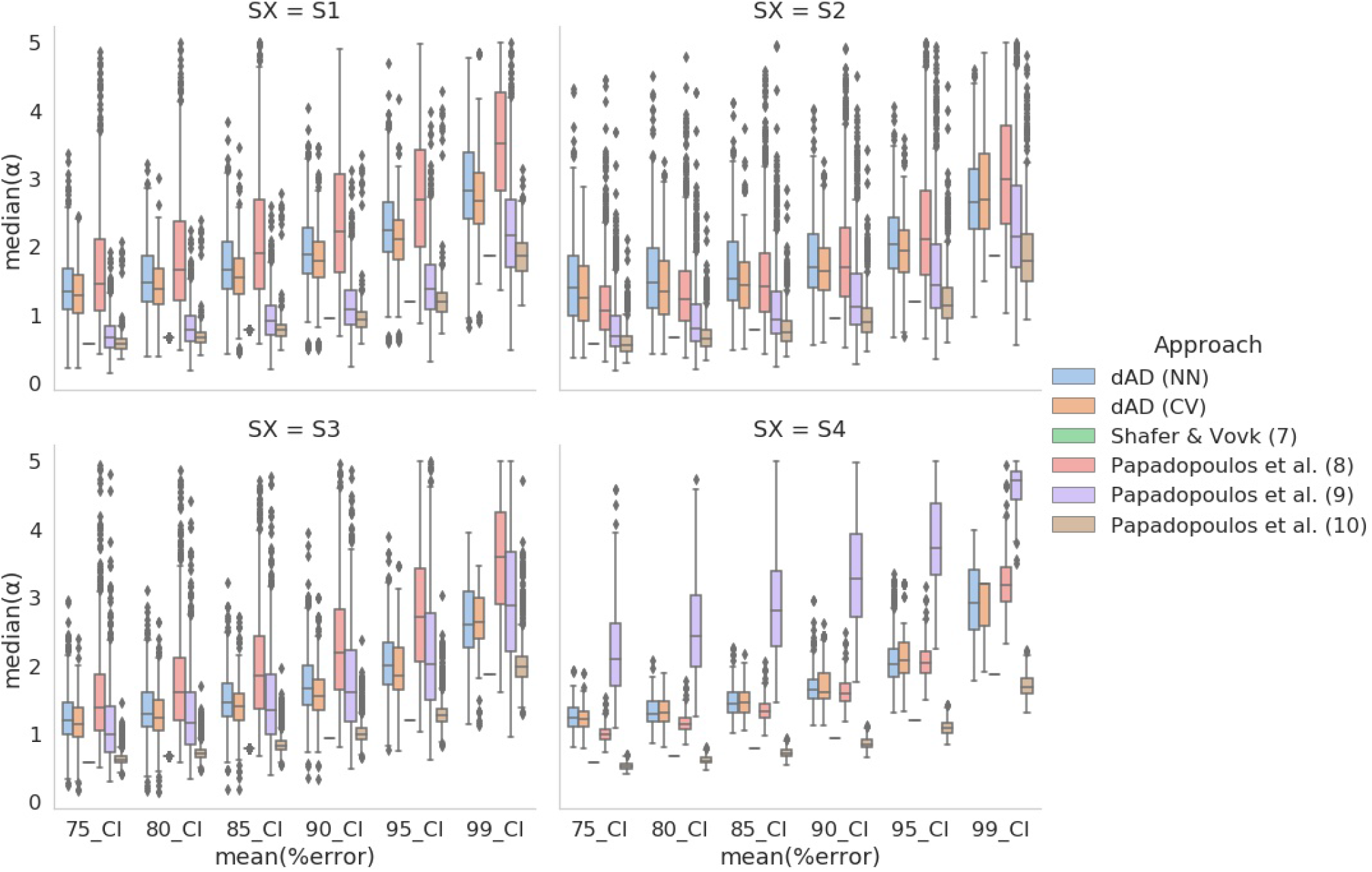
Comparison of original study by Shafer and Vovk [2008], normalization approach with underlying error model Papadopoulos and Haralambous [2010], normalization approach with normalized standard deviation Papadopoulos et al. [2011] and normalized distances Papadopoulos et al. [2011], and two proposed dAD approaches with nonconformity scores calculated as the difference from the mean (NN) or difference from the cross-validation prediction (CV). All five of them are compared over SCKBA dataset with four testing scenarios (S1-S4).

**Figure 5:**
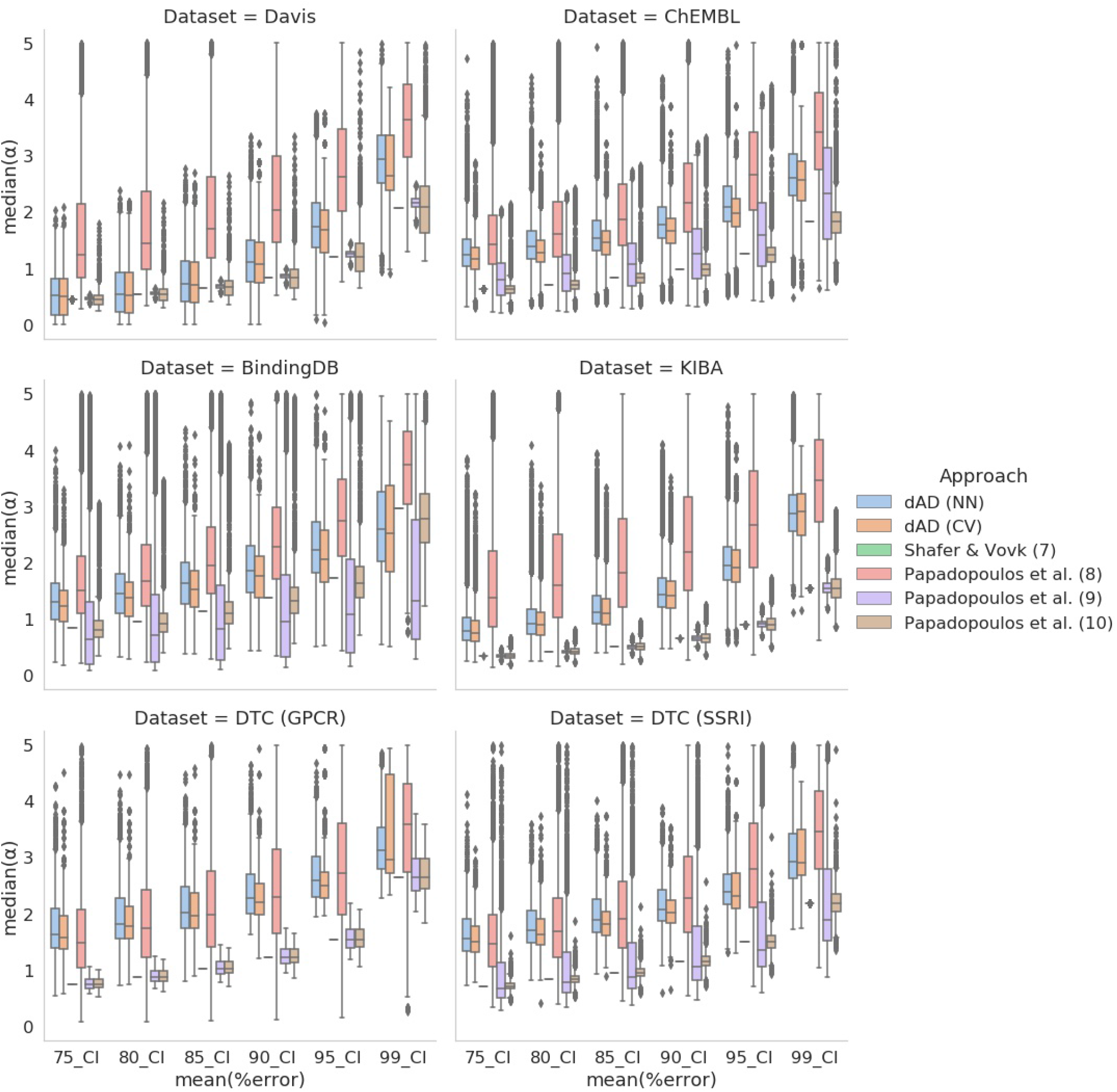
Comparison of original Shafer and Vovk study Shafer and Vovk [2008], normalization approach with underlying error model Papadopoulos and Haralambous [2010], normalization approach with normalized standard deviation Papadopoulos et al. [2011] and normalized distances Papadopoulos et al. [2011], and two proposed dAD approaches with nonconformity scores calculated as the difference from the mean (NN) or difference from the cross-validation prediction (CV). All five of them are compared over six benchmark datasets.

## 6 Summary

The role of Applicability Domain and Conformal Predictors framework in Drug discovery context is to provide improved machine learning model performance assessment on the level of individual predictions. This work merges ideas of an applicability domain (AD) and conformal predictors (CP) to provide improved guarantees in bioactivity type of tasks, involving complex dyadic data spaces such as compound-target binding affinity prediction. As stated in Section 2, classic CP frameworks rely on fixed calibration sets which are based on their overall distributional similarity to the training data. Compound-target binding affinity problem is defined by independently distributed chemical and biological spaces and models produced over such datasets may exhibit significantly different performance over different clusters of compounds and target families dependent on their distribution in the training set and model properties. Moreover, realistic use-cases of such machine learned models in drug discovery or drug repurposing, ask for predictions over novel compounds or targets for which exchangeability requirement is only marginally satisfied. This was a motivation for developing a dynamic conformal predictor framework that involves a design of dynamic calibration sets - sets that are defined by the training set samples neighbouring the tested sample. The primary concern in the context of developing of the new framework was the effective treatment of the dyadic character of the data: i.e. input data involves interaction of two types of entities and their own distributions.

We solve this duality problem in defining conformity regions, separately for each tested sample, by searching for its neighbours, independently in the compound and protein target space, and retrieving experimentally measured binding affinities for the compounds and targets residing in this neighbourhood. Consequently, this method produces prediction regions that are specific for an individual test sample given the confidence level of interest, without the need to normalise resulting nonconformity scores. We prove in this work that dynamic calibration sets more accurately reflect the performance of the model in the area close to the tested sample, providing more robust guarantees for suggested prediction regions. Our experiments on the compound-target binding problem prove that this approach provides similar performance in terms of validity and prediction regions as other *state-of-the-art* CP approaches based on normalised nonconformity scores Papadopoulos and Haralambous [2010] Papadopoulos et al. [2011] for the standard testing scenario (S1). However, for more difficult scenarios, such as drug or target discovery and drug repurposing (S2-S4), involving test samples in or out of the borders of the training data space, proposed dAD approach proved to be more effective, providing validity over reasonable prediction regions, which proved not to be the case with other CP approaches.

## References

Anna Cichonska, Balaguru Ravikumar, Elina Parri, Sanna Timonen, Tapio Pahikkala, Antti Airola, Krister Wennerberg, Juho Rousu, and Tero Aittokallio. Computational-experimental approach to drug-target interaction mapping: a case study on kinase inhibitors. PLoS computational biology, 13(8):e1005678, 2017.

Anna Cichońska, Balaguru Ravikumar, Robert J Allaway, Fangping Wan, Sungjoon Park, Olexandr Isayev, Shuya Li, Michael Mason, Andrew Lamb, Ziaurrehman Tanoli, et al. Crowdsourced mapping of unexplored target space of kinase inhibitors. Nature communications, 12(1):1–18, 2021.

Hakime Öztürk, Arzucan Özgür, and Elif Ozkirimli. Deepdta: deep drug–target binding affinity prediction. Bioinformatics, 34(17):i821–i829, 2018.

Thin Nguyen, Hang Le, Thomas P Quinn, Tri Nguyen, Thuc Duy Le, and Svetha Venkatesh. Graphdta: Predicting drug–target binding affinity with graph neural networks. Bioinformatics, 37(8):1140–1147, 2021.

Sangsoo Lim, Yijingxiu Lu, Chang Yun Cho, Inyoung Sung, Jungwoo Kim, Youngkuk Kim, Sungjoon Park, and Sun Kim. A review on compound-protein interaction prediction methods: data, format, representation and model. Computational and Structural Biotechnology Journal, 19:1541–1556, 2021.

Xing Chen, Chenggang Clarence Yan, Xiaotian Zhang, Xu Zhang, Feng Dai, Jian Yin, and Yongdong Zhang. Drug– target interaction prediction: databases, web servers and computational models. Briefings in bioinformatics, 17(4): 696–712, 2016.

Thomas N Kipf and Max Welling. Semi-supervised classification with graph convolutional networks. arXiv preprint arXiv:1609.02907, 2016.

Bharath Ramsundar, Peter Eastman, Patrick Walters, and Vijay Pande. Deep learning for the life sciences: applying deep learning to genomics, microscopy, drug discovery, and more. O’Reilly Media, 2019.

Domenico Gadaleta, Giuseppe Felice Mangiatordi, Marco Catto, Angelo Carotti, and Orazio Nicolotti. Applicability domain for qsar models: where theory meets reality. International Journal of Quantitative Structure-Property Relationships (IJQSPR), 1(1):45–63, 2016.

Natália Aniceto, Alex A Freitas, Andreas Bender, and Taravat Ghafourian. A novel applicability domain technique for mapping predictive reliability across the chemical space of a qsar: reliability-density neighbourhood. Journal of cheminformatics, 8(1):1–20, 2016.

Waldemar Klingspohn, Miriam Mathea, Antonius Ter Laak, Nikolaus Heinrich, and Knut Baumann. Efficiency of different measures for defining the applicability domain of classification models. Journal of cheminformatics, 9(1): 1–17, 2017.

Miriam Mathea, Waldemar Klingspohn, and Knut Baumann. Chemoinformatic classification methods and their applicability domain. Molecular Informatics, 35(5):160–180, 2016.

Alex Gammerman, Volodya Vovk, and Vladimir Vapnik. Learning by transduction. arXiv preprint arXiv:1301.7375, 2013.

Glenn Shafer and Vladimir Vovk. A tutorial on conformal prediction. Journal of Machine Learning Research, 9(3), 2008.

Harris Papadopoulos. Inductive conformal prediction: Theory and application to neural networks. INTECH Open Access Publisher Rijeka, 2008.

Ulf Johansson, Henrik Boström, Tuve Löfström, and Henrik Linusson. Regression conformal prediction with random forests. Machine learning, 97(1):155–176, 2014.

Harris Papadopoulos and Haris Haralambous. Neural networks regression inductive conformal predictor and its application to total electron content prediction. In International Conference on Artificial Neural Networks, pages 32–41. Springer, 2010.

Harris Papadopoulos, Vladimir Vovk, and Alexander Gammerman. Regression conformal prediction with nearest neighbours. Journal of Artificial Intelligence Research, 40:815–840, 2011.

Jonathan Alvarsson, Staffan Arvidsson McShane, Ulf Norinder, and Ola Spjuth. Predicting with confidence: using conformal prediction in drug discovery. Journal of Pharmaceutical Sciences, 110(1):42–49, 2021.

Vladimir Vovk, Ilia Nouretdinov, Valery Manokhin, and Alexander Gammerman. Cross-conformal predictive distributions. In Conformal and Probabilistic Prediction and Applications, pages 37–51. PMLR, 2018.

Jing Tang, Balaguru Ravikumar, Zaid Alam, Anni Rebane, Markus Vähä-Koskela, Gopal Peddinti, Arjan J van Adrichem, Janica Wakkinen, Alok Jaiswal, Ella Karjalainen, et al. Drug target commons: a community effort to build a consensus knowledge base for drug-target interactions. Cell chemical biology, 25(2):224–229, 2018.

Mindy I Davis, Jeremy P Hunt, Sanna Herrgard, Pietro Ciceri, Lisa M Wodicka, Gabriel Pallares, Michael Hocker, Daniel K Treiber, and Patrick P Zarrinkar. Comprehensive analysis of kinase inhibitor selectivity. Nature biotechnology, 29(11):1046–1051, 2011.

Jing Tang, Agnieszka Szwajda, Sushil Shakyawar, Tao Xu, Petteri Hintsanen, Krister Wennerberg, and Tero Aittokallio. Making sense of large-scale kinase inhibitor bioactivity data sets: a comparative and integrative analysis. Journal of Chemical Information and Modeling, 54(3):735–743, 2014.

Michael K Gilson, Tiqing Liu, Michael Baitaluk, George Nicola, Linda Hwang, and Jenny Chong. Bindingdb in 2015: a public database for medicinal chemistry, computational chemistry and systems pharmacology. Nucleic acids research, 44(D1):D1045–D1053, 2016.

Anna Gaulton, Louisa J Bellis, A Patricia Bento, Jon Chambers, Mark Davies, Anne Hersey, Yvonne Light, Shaun McGlinchey, David Michalovich, Bissan Al-Lazikani, et al. Chembl: a large-scale bioactivity database for drug discovery. Nucleic acids research, 40(D1):D1100–D1107, 2012.

Greg Landrum, Paolo Tosco, Brian Kelley, sriniker, gedeck, NadineSchneider, Riccardo Vianello Ric, Andrew Dalke, Brian Cole, and et al. rdkit/rdkit: 2019093(q32019)release. Jan2020. doi : 10.5281/zenodo.3603542.

Adam Paszke, Sam Gross, Francisco Massa, Adam Lerer, James Bradbury, Gregory Chanan, Trevor Killeen, Zeming Lin, Natalia Gimelshein, Luca Antiga, et al. Pytorch: An imperative style, high-performance deep learning library. Advances in neural information processing systems, 32, 2019.

Yixuan He, Xitong Zhang, Junjie Huang, Mihai Cucuringu, and Gesine Reinert. Pytorch geometric signed directed: A survey and software on graph neural networks for signed and directed graphs. arXiv preprint arXiv:2202.10793, 2022.

Jeffrey Jie-Lou Liao. Molecular recognition of protein kinase binding pockets for design of potent and selective kinase inhibitors. Journal of medicinal chemistry, 50(3):409–424, 2007.

Robert Roskoski Jr. A historical overview of protein kinases and their targeted small molecule inhibitors. Pharmacological research, 100:1–23, 2015.

R Roskoski Jr. Enzyme structure and function. 2014.

Nan Xiao, Dong-Sheng Cao, Min-Feng Zhu, and Qing-Song. Xu. protr/ProtrWeb: R package and web server for generating various numerical representation schemes of protein sequences. Bioinformatics, 31(11):1857–1859, 2015. ISSN 1367-4803. doi:10.1093/bioinformatics/btv042. URL http://bioinformatics.oxfordjournals.org/content/31/11/1857.

Tianqi Chen and Carlos Guestrin. Xgboost: A scalable tree boosting system. In Proceedings of the 22nd acm sigkdd international conference on knowledge discovery and data mining, pages 785–794, 2016.

