## Supplementary figures and images for "Dynamic applicability domain (*d*AD) for compound-target binding affinity prediction task with confidence guarantees"

### Figure 1, Supplementary

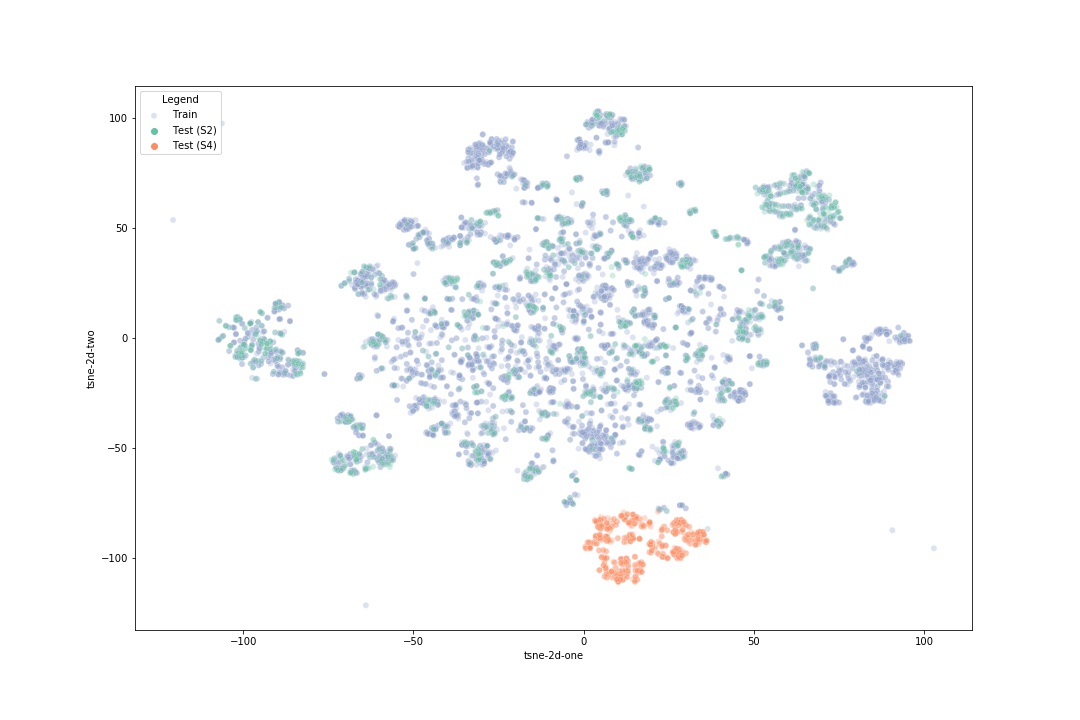

### Figure 2, Supplementary

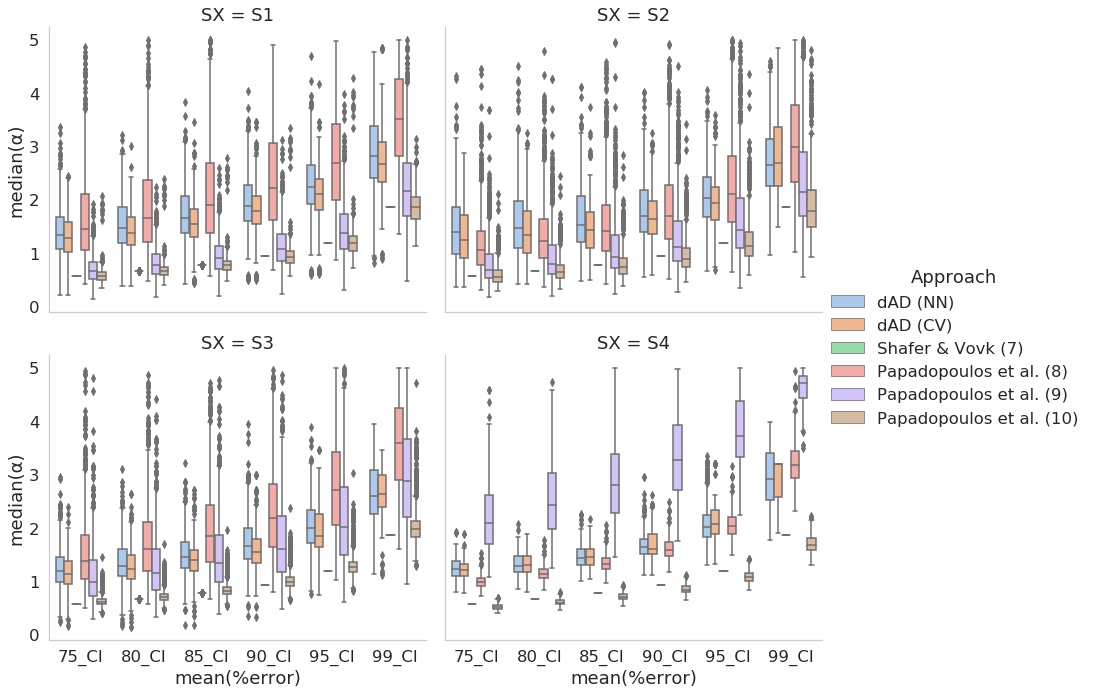

### Figure 3, Supplementary

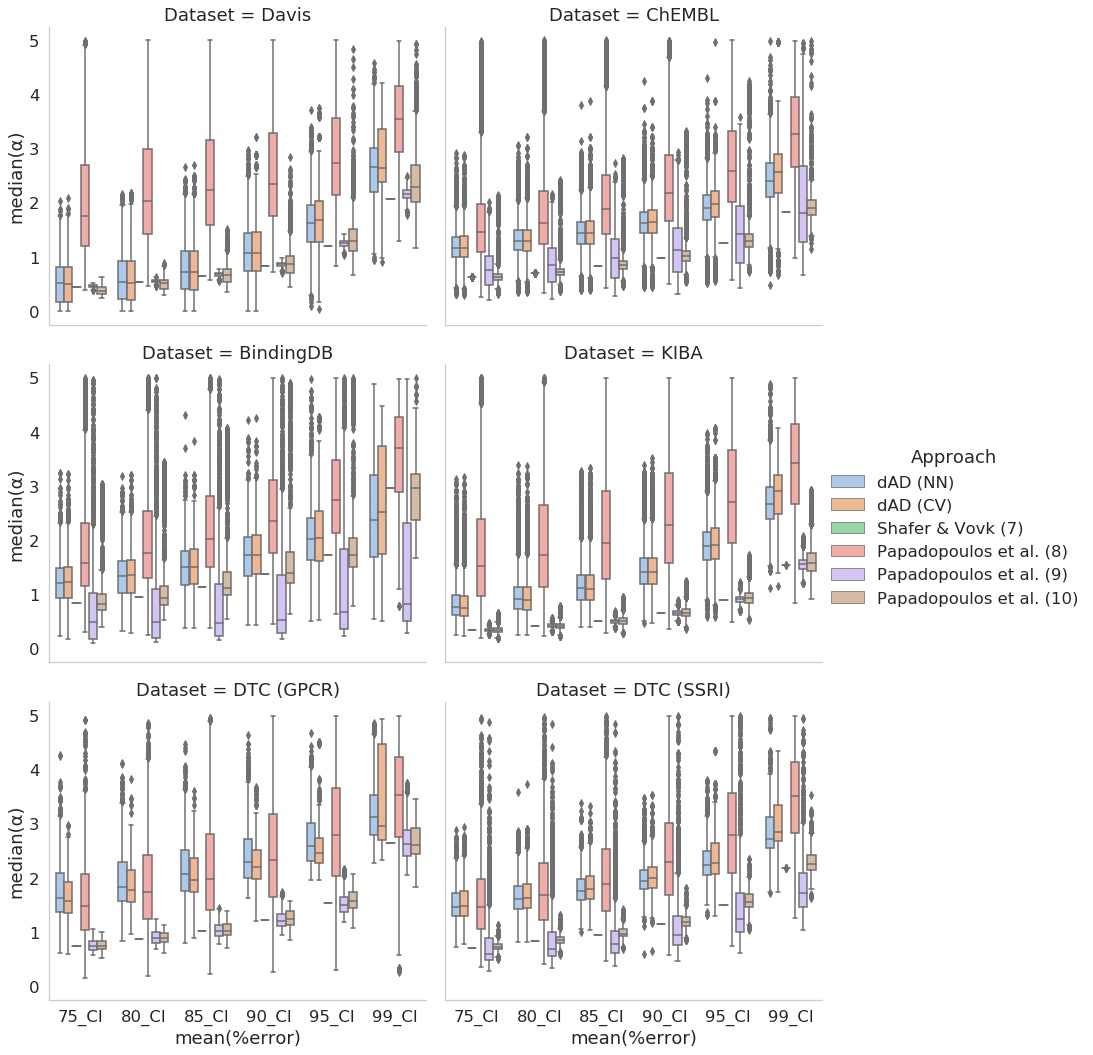

### Figure 4, Supplementary

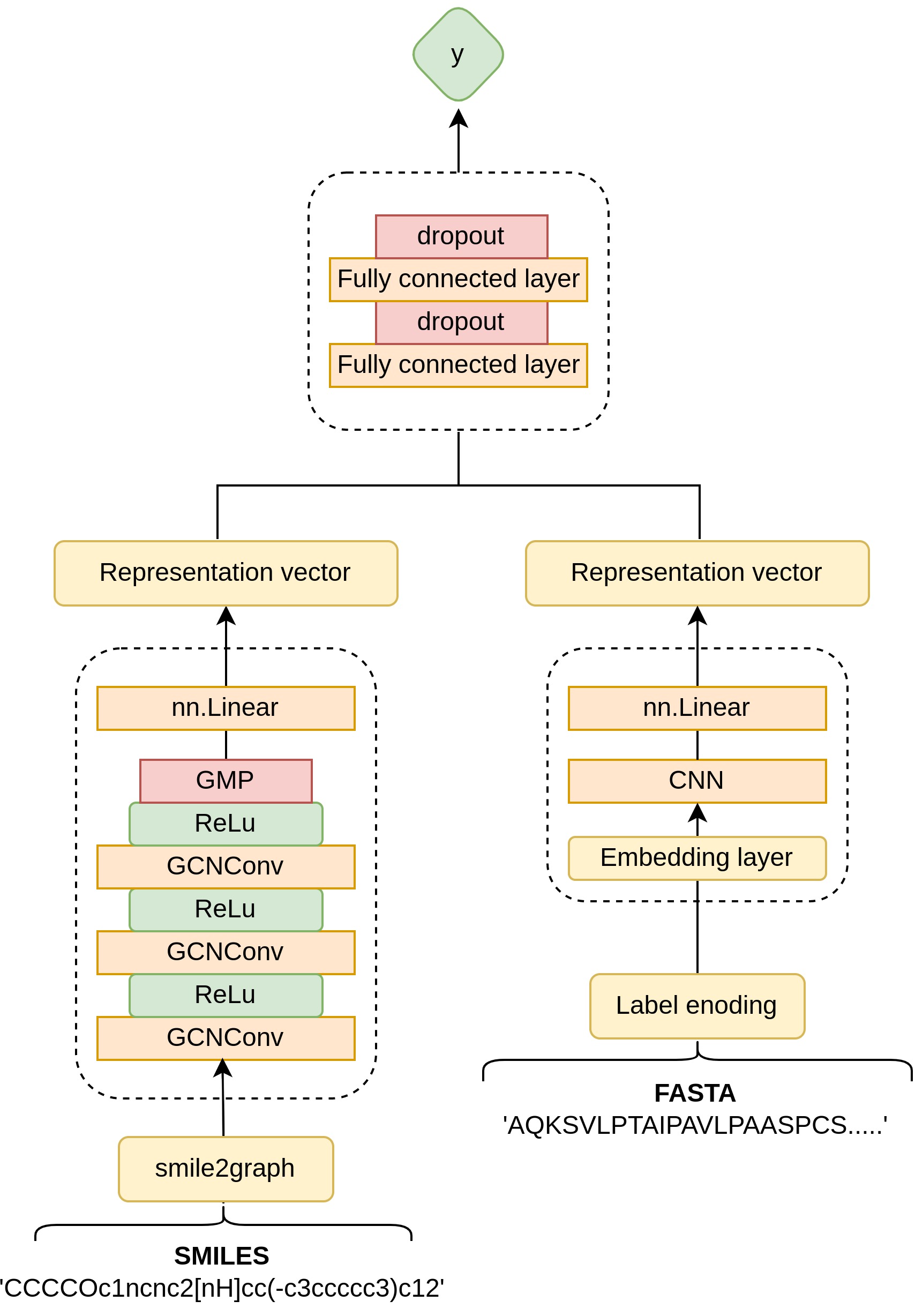

### Figure 5, Supplementary

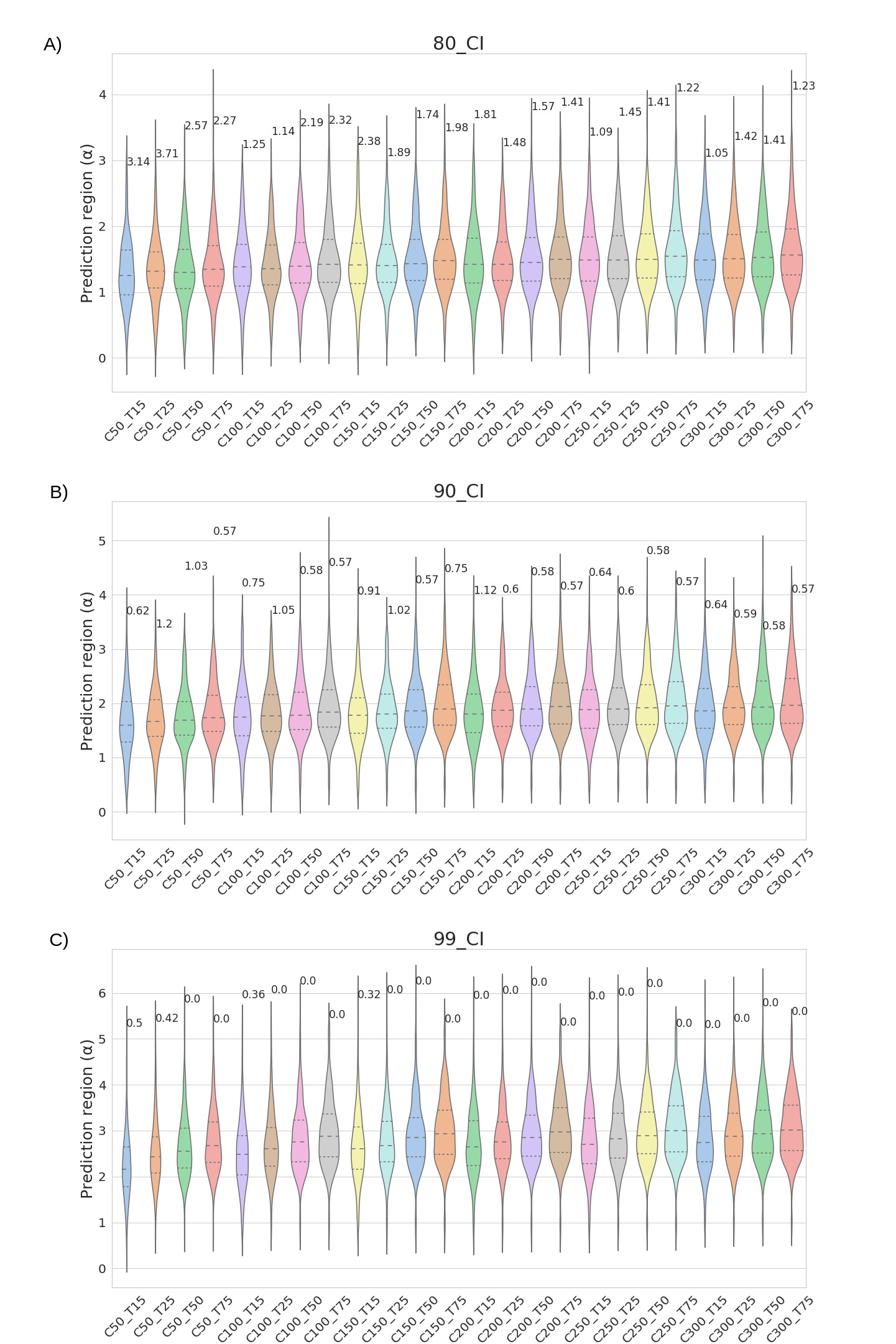

### Figure 6, Supplementary

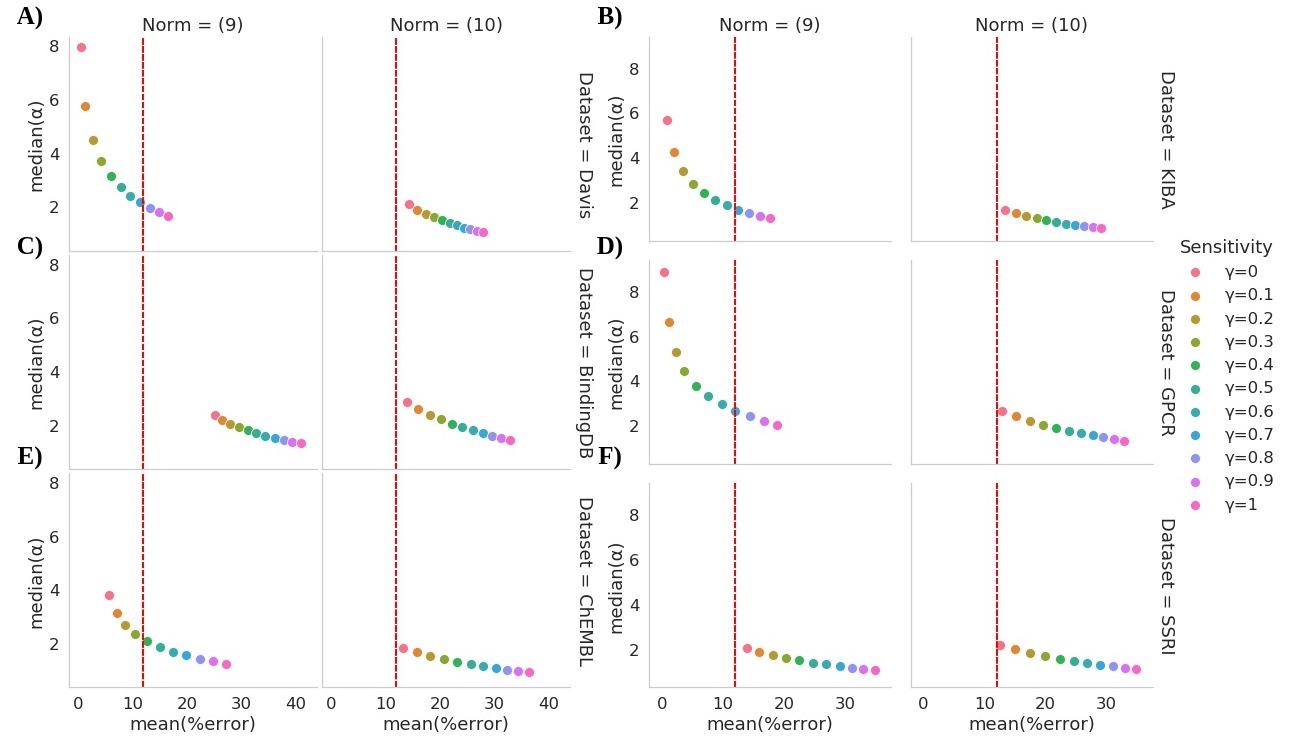

### Figure 7, Supplementary

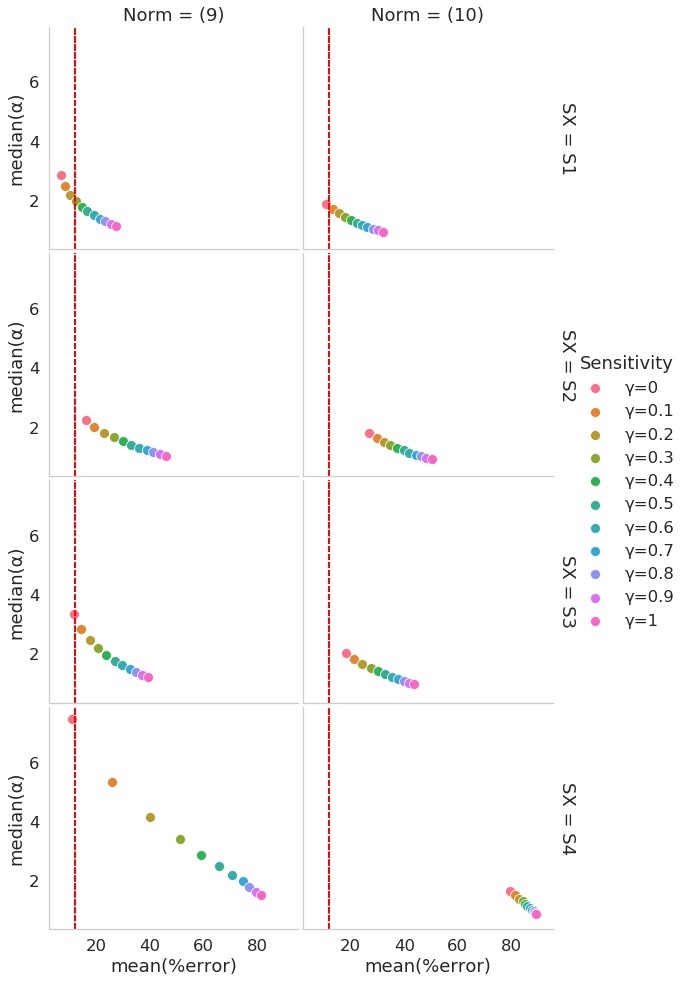

### Figure 8, Supplementary

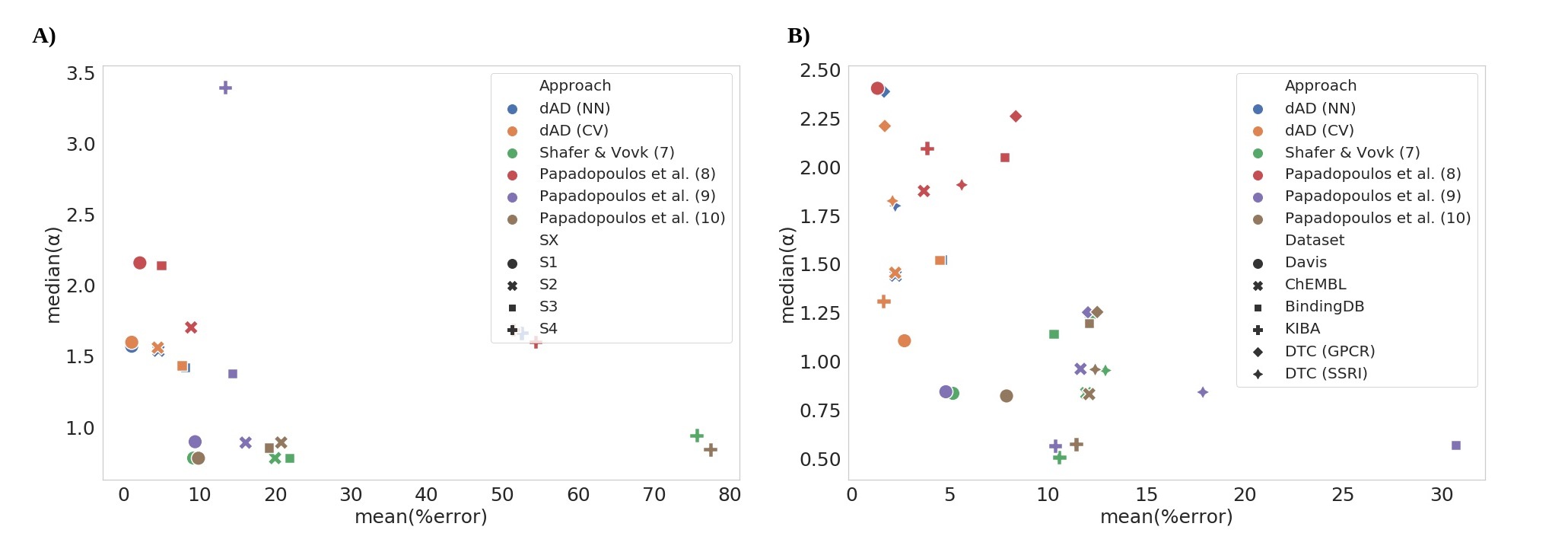
